# Integrated sequencing approach to probe rRNA modification landscape during human embryonic stem cell differentiation

**DOI:** 10.64898/2026.08.10.743918

**Authors:** Tessa W. Y. Chan, Ivana Barbaric, Emma E. Thomson

**Affiliations:** School of Biosciences, The University of Sheffield, Sheffield, S10 2TN, UK

## Abstract

The ribosome, long regarded as a passive, uniform machine, has only recently been recognised as a direct regulator of translation. Mass spectrometry and sequencing approaches have shown that heterogeneity in ribosome composition exists, which can actively regulate the translational process. One source of this heterogeneity is the modification of ribosomal RNA (rRNA), primarily pseudouridylation (pseU) and 2′-*O*-methylation (2OMe), mediated by specific H/ACA and C/D box small nucleolar RNAs (snoRNAs). Here, we investigate how the stoichiometry of rRNA modifications varies during embryonic stem cell differentiation. Using the modification basecalling capability of Nanopore direct RNA sequencing, we have identified distinct stoichiometric changes in modification patterns between pluripotent and differentiated cells, revealing highly dynamic, site-specific regulation. Further, profiling of snoRNA expression during trilineage differentiation revealed differential expression of H/ACA and C/D box snoRNAs responsible for a subset of these dynamic modifications. By integrating rRNA and snoRNA sequencing approaches, we have built a comprehensive profile of rRNA modification dynamics during early embryonic cell fate decisions, highlighting potential regulatory mechanisms for ribosome heterogeneity during development.

**GRAPHICAL ABSTRACT:** 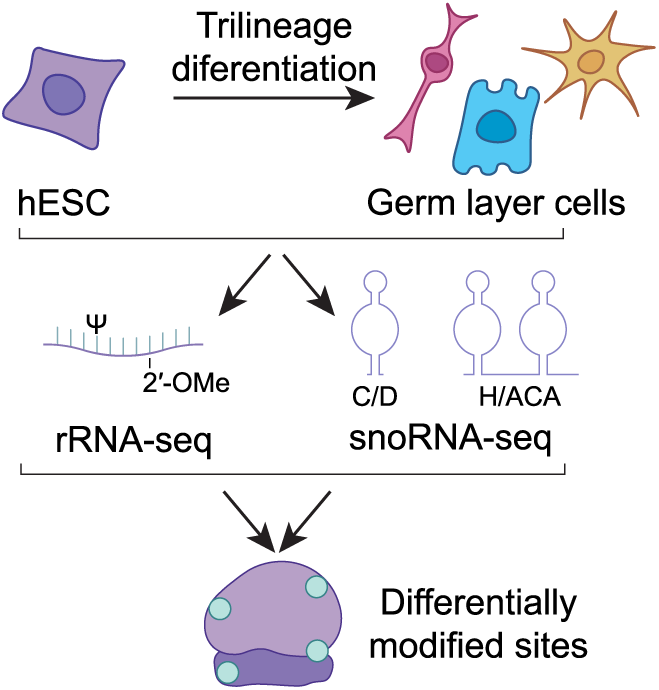

## INTRODUCTION

Ribosomes are the ancient macromolecular ribonucleoprotein particles responsible for carrying out the universally conserved process of protein synthesis. In humans, the small (40S) and large (60S) ribosomal subunits comprise four rRNA species (18S, 5.8S, 28S and 5S) and 80 ribosomal proteins (r-proteins), both of which are extensively modified. The two most prevalent rRNA modifications are 2′-*O-*methylation (2OMe) and pseudouridylation (pseU) [1,2,3], which are predominantly introduced co-transcriptionally by C/D box and H/ACA box small nucleolar ribonucleoproteins (snoRNPs), respectively [4,5,6]. Human guide snoRNA-rRNA interactions display complex targeting dynamics, including one-to-one, one-to-many, and many-to-one [7,8]. Consequently, because of this lack of a strict one-to-one relationship, expression levels of individual snoRNAs are not necessarily indicative of the modification stoichiometries of the rRNA targets [9,10]. However, a subset of rRNA sites that exhibit dynamic changes in modification stoichiometry demonstrate a correlation with the expression of their specific guide snoRNAs [10,11,12].

While translation is a highly regulated process, until recently, the ribosome itself had not been shown to exhibit any intrinsic regulatory capacity. Rather, investigations of translation regulation have largely focused on the role of translation factors and transcript features [13,14,15]. However, this paradigm has shifted following the discovery that compositional heterogeneity within the ribosome can drive downstream effects on translation [16,17]. The extent of heterogeneity in the composition of ribosomes has only recently begun to be appreciated, with advances in mass-spec and sequencing methodologies revealing variation in both r-proteins and rRNA composition. Variation in r-protein composition has been seen in a number of different contexts including differential paralogue incorporation [18,19,20], changes in protein stoichiometry [21,22,23] and post-translational modifications [23,24]. In the case of the rRNA, primary sequence variants [25,26,27,28] in addition to changes in modification stoichiometry have been reported [9,11,12,29,30,31]. Notably, compositional variation that occurs in different biological contexts has been observed between different cell types (intercellular heterogeneity), within a cell type (intracellular heterogeneity) and in response to developmental and environmental cues [16,17]. While these changes in ribosome composition do not by default impart a functional specialisation, a growing number of studies propose mechanisms through which compositional change leads to a change in translational output [11,12,19,22].

Changes in ribosome composition have been identified during crucial cell fate decisions where pluripotent embryonic stem cells (ESC) differentiate into lineage committed populations. This has led to the suggestion that ribosome heterogeneity contributes to the rapid reprogramming of translation during early differentiation [12,22]. The role of rRNA modification heterogeneity during cell fate decisions has been explored for 2OMe and has implicated changes in modification patterns of specific residues in the regulation of gene expression during neural development and early stem cell commitment [12]. However, a comprehensive profile of rRNA pseudouridylation in stem cells and how stoichiometry changes during differentiation remains uncharacterized.

Historically, sequencing studies investigating pseU (such as CMC-seq and HydraPsi-seq) [32,33] and 2OMe (RiboMeth-seq) [34,35] in rRNA have relied on next generation sequencing (NGS) coupled with chemical treatments of RNAs. While informative, these approaches rely on inferring the modification status of residues rather than detecting the native modification directly [36]. Similarly, while early versions of third-generation long read direct RNA sequencing (DRS) detected each of the canonical RNA residues, modification was inferred through the error rates of individual bases [37,38], or detected through custom modification basecalling models [39,40,41]. Recently, however, advances in basecalling models have enabled the direct, simultaneous detection of native modifications, including the 2OMe and pseU, within single RNA molecules.

Here we employ DRS to determine the stoichiometries of all 2OMe and pseU residues across each rRNA species in human ESCs (hESC) and determine how these change during lineage commitment into the three germ layers, which mimic gastrulation as a significant change in cellular potency. Further, by pairing this direct modification profiling with next generation sequencing of snoRNAs, we demonstrate that snoRNA expression correlates with changes in stoichiometry for a subset of modified sites. These findings provide a single-nucleotide view on rRNA modification dynamics, demonstrating that specific layers of ribosome heterogeneity are precisely regulated during early human cell fate decisions.

## MATERIAL AND METHODS

### Tissue culture

#### Cell growth and maintenance

Human embryonic stem cell line MasterShef11 (Mshef11; RRID:CVCL_BW83) was maintained in Essential 8 (E8; Gibco) or TeSR-E8 media (STEMCELL Technologies) on vitronectin (Gibco). Media changes were performed daily. Cells were passaged using ReLeSR™ (STEMCELL Technologies) when reaching 80% confluency, and split at a ratio of 1:3-1:6. Cells were harvested by single cell dissociation using 1X TrypLE™ Express (Gibco).

Cells were thawed in E8 supplemented with 10μM Y-27632 (ROCKi; STEMCELL Technologies). Cells were cryopreserved in 10% v/v DMSO (Sigma-Aldrich) in FBS (BIOSERA) or in STEM- CELLBANKER® (AMSBIO).

Copy number variation of the working cell bank was assessed using Next generation low-pass sequencing with all cells used being verified as being karyotypically normal. Experiments were carried out from passage 3 onwards, post-thawing.

### Trilineage differentiation

MShef11 was differentiated into the three primary germ layers using STEMdiff™ Trilineage Differentiation Kit (STEMCELL Technologies) in a 6-well plate format, following the manufacturer’s instructions. In brief, cells were dissociated into single cells using Gentle Cell Dissociation Reagent (STEMCELL Technologies). At day 0, cells were suspended in 10μM ROCKi-containing E8 media, and seeded at 2×10^6^ cells/well for ectoderm and endoderm differentiation, or 0.5×10^6^ cells/well for mesoderm differentiation. On day 1, plating media was replaced with the corresponding lineage differentiation media. Media was replaced every day, until day 5 when mesoderm and endoderm were harvested, or until day 7 when ectoderm was harvested. Differentiation was confirmed by RT-qPCR using pluripotent and lineage specific markers.

### RNA analysis

#### RNA extraction

Total RNA was extracted using TRI Reagent™ (Sigma-Aldrich) and chloroform (Fisher Chemical) following the manufacturer’s instructions. To remove DNA contaminants, RNA pellets were dissolved in a 50μL DNase reaction made from 1X TURBO DNase™ buffer, and 2U of TURBO DNase™ (Invitrogen). DNase reactions were incubated for 30min at 37°C. Following DNase treatment, RNA samples were purified using Monarch® Spin RNA Cleanup Kit (New England Biolabs) according to the manufacturer’s instructions.

RNA concentration was measured using Qubit™ RNA Broad Range Assay Kit with the Qubit 4 Fluorometer (Invitrogen) and RNA integrity was assessed using RNA ScreenTape with the 4150 TapeStation System (Agilent).

### RT-qPCR

Reverse transcription was carried out with random hexamers using TaqMan™ Reverse Transcription Reagents (Invitrogen) following the manufacturer’s instructions. 1μg of RNA was reverse transcribed per reaction. cDNA was diluted 10-fold before RT-qPCR.

qPCR reactions were prepared as 10μL reactions containing 2X PowerUp™ SYBR™ Green Master Mix (Applied Biosystems), 0.3μM forward primer, 0.3μM reverse primer, and 5ng of cDNA. qPCR was conducted in QuantStudio 12K Flex (Applied Biosystems) using a 384-well format and in standard mode. qPCR data analysis was carried out using the standard curve method.

Primers for pluripotency and lineage-specific markers are summarised in Table S1.

### In vitro transcription (IVT)

pcDNA3.1 plasmids containing human 5S, 5.8S, 18S, and 28S rRNA under a T7 promoter were linearised by restriction digest. 100pmol (5S, 5.8S) or 500pmol (18S, 28S) of linearised plasmids were transcribed in a 100μL reaction containing 12mM MgCl2, 40mM Tris pH8.0, 2mM spermidine, 10mM NaCl, 0.01% v/v Triton X-100, 3.6mM of each NTP (Thermo Scientific), 2U pyrophosphatase (NEB), 40U RNaseOUT™ (Invitrogen), and 49.1nM T7 RNA polymerase (P266L mutant, made in-house). The reaction was incubated at 37°C for 4h. DNase treatment was carried out using 2U of TURBO DNase, at 37°C for 30min. RNA was purified using Monarch® Spin RNA Cleanup Kit.

### Nanopore direct RNA sequencing

Total RNA from MShef11 hESCs, trilineage differentiated cells, and IVT rRNA was used as input for preparing Nanopore direct RNA sequencing libraries. For DRS, three biological replicates of each cell type, and two replicates of IVT were used. Library preparation was carried out using Direct RNA-sequencing kit (Oxford Nanopore Technologies #SQK-RNA004). 1μg of total RNA was purified using 1.8X volume of RNAClean XP beads (Beckman Coulter) and eluted in 16μL of water.

The purified RNA was polyadenylated by adding 1X Poly(A) Polymerase Reaction buffer, 1mM ATP, 20U of murine RNase inhibitor (NEB), and 5U of *E. coli* poly(A) polymerase (NEB), made up to a 20μL reaction. The reaction mixture was incubated at 37°C for 30s, then quenched by adding EDTA to a final concentration of 10mM. The polyadenylated RNA was purified using 1.8X RNAClean XP beads, and eluted in 6.5μL of water. For IVT samples, 500ng of each rRNA were polyadenylated, purified, and eluted in 12μL of water. The polyadenylated IVT rRNAs were mixed at a molar ratio of 10:10:30:50fmol (5S:5.8S:18S:28S).

300ng of polyadenylated RNA, or 100ng of mixed IVT rRNA were ligated with sequencing adaptors in a 7.5μL reaction, by mixing RNA with 20U of murine RNase inhibitor, 0.093μM of annealed RT adapters, 1X of NEBNext® Quick Ligation Reaction Buffer (NEB), and 1500U of T4 DNA ligase (NEB). The ligation reaction was incubated for 5min at room temperature. Custom four-sample SeqTagger barcodes were used [42] (Table S1).

Adapter ligated RNA was reverse transcribed in a 20μL reaction with 0.5mM of dNTP, 1X Induro RT Reaction Buffer, and 200U of Induro® Reverse Transcriptase (NEB). Induro RT reaction was incubated at 60°C for 30min, then inactivated at 70°C for 10min.

Following RT, the RNA:cDNA library was purified using 1.8X volume of RNAClean XP beads, and eluted in 12μL of water. Library concentration was quantified using Qubit 1X dsDNA High Sensitivity Assay Kit (Invitrogen), and library size distribution was assessed using TapeStation RNA ScreenTape. Four samples were multiplexed in a single sequencing run with equal ng of library from each sample made up to 23μL with water. The RNA:cDNA library was ligated to sequencing motor protein and purified following manufacturer’s instructions.

Libraries were sequenced using a MinION RNA flow cell (Oxford Nanopore Technologies #FLO-MIN004RA) and a MinION Mk1B or Mk1D device. Sequencing libraries and the flow cells were primed prior to sequencing following manufacturer’s instructions, and the libraries were sequenced for a maximum of 2.0Gb, or up to 12h in sequencing time.

After each sequencing run, the flow cell was washed using the Flow Cell Wash Kit (Oxford Nanopore Technologies). The wash mix was composed of 2μL of DNase-containing Wash Mix WMX, 1μL of RNase H (NEB), 1μL of RNase Cocktail™ Enzyme Mix (Invitrogen), and 396μL of Wash Diluent DIL.

### Illumina small RNA sequencing

500ng of RNA from MShef11 hESCs and trilineage differentiated cells was used as input for preparing small RNA-seq libraries, using a Small RNA-Seq Library Prep Kit (Lexogen) according to the manufacturer’s instructions, with 12 cycles of PCR carried out. After library preparation, the concentration of each library was quantified using Qubit dsDNA High Sensitivity Quantification Assay Kit (Invitrogen). cDNA size distribution for each library was assessed using High Sensitivity D1000 ScreenTape (Agilent).

A size selection was carried out to remove <100bp DNA fragments, which corresponded to sequencing library linkers. 9μL of each sequencing library was incubated with 1.8X volumes of AMPure XP Beads (Beckman Coulter) for 5min at room temperature. The mixture was immobilized on a magnetic rack and the supernatant removed. The beads were washed twice with 70% ethanol, and eluted in 8μL of elution buffer provided in the library preparation kit. Library concentration was quantified using Qubit and size selection was assessed using TapeStation. Using the TapeStation Analysis software, the average library size was calculated by drawing a 100-1000bp range for each library. The molarity of each library was then estimated using the average library size and the DNA concentration assessed by Qubit. 30fmol of each library was multiplexed into a final volume of 25μL of elution buffer. The multiplexed library was sequenced using paired-end 150bp NovaSeq™ X Plus (Illumina) for a minimum of 375Gb by Genewiz.

### Data Analysis

Sequencing analysis was carried out using high performance computing. The scripts used for data analysis are deposited in Github (https://github.com/Thomson-RNA-Lab/Integrated-rRNA-snoRNA-seq). Data visualisation was carried out using R.

### Nanopore direct RNA sequencing

Raw reads as FAST5 files were demultiplexed using SeqTagger v1.0d [42]. The resulting demultiplexed FAST5 files were then converted to POD5. POD5 reads were basecalled using dorado v1.3.0 (https://github.com/nanoporetech/dorado). Reads were basecalled using super-accurate mode with the basecalling model rna004_130bps_sup@v5.2.0, with modification information for all available RNA modification types, and a minimum quality score threshold of 8: *basecaller sup*, *pseU*_2*OmeU*, *m*5*C*_2*OmeC*, *inosine*_*m*6*A*_2*OmeA*, 2*OmeG* − −*min* −*qscore* 8. Finally, the basecalled BAM files were converted to FASTQ using samtools with the options −*T* “ ∗ ”, which retained all modified basecall information during the conversion.

Sequencing reads were aligned to a reference transcriptome, composed of the human 18S (NR_145820.1), 28S (NR_003287.4), 5.8S (NR_003285.3) and 5S (NR_023363.1). FASTQ reads were mapped with minimap2 [43] with the options −*ayx map* − *ont* − *k*14 − −*cs* = *long* −*w*5 − *N* 10 − −*secondary* = *no*, resulting in an aligned BAM file. Sequencing alignment for each sample was assessed by NanoCount [44] and nanoq [45]. Output files of NanoCount and nanoq were then summarised using mutiqc [46].

Modification information from the read alignment was analysed using modkit v0.5.0 (https://github.com/nanoporetech/modkit). Using the aligned reads of the merged IVT samples, 20% of aligned reads were subsampled to acquire the probability distribution of modified basecalls (*sample* − *probs* − −*mapped* − *only* − −*include* − *bed* < *ground*_*truth*. *bed* > − − *region* < 18*S*/28*S*/5.8*S* > − − *sampling* − *frac* 0.2). An optimal probability threshold was determined such that a 99.5% precision in modification basecalling of the merged IVT samples was achieved. The probability thresholds were then applied to all other samples to remove low confidence basecalls during pileup (*pileup* − −*region* < 18*S*/28*S*/5.8*S* > − − *max* − *depth* 100000 − −*filter* − *threshold* < *A*/*C*/*G*/*T* > <*optimal threshold* >). Subsequently, the modification percentage at each position of the reference rRNA sequence was calculated by counting the number of modified basecalls (*N_mod_*) over the total number of valid basecalls per position (*N_valid_*), where *N_valid_* = *N_mod_* + *N_canonical_* . The quantification files were filtered for all rRNA positions annotated to be pseU and 2OMe sites.

A generalised linear model was made to model modification level. For each replicate sample, the *N_mod_* for position *i* in cell type *j* was modelled to follow a negative binomial distribution, dependent on position *i*, cell type *j* and the interaction between the two variables, offset by *N_valid_* of the corresponding sample. For each position, the pairwise difference in estimated 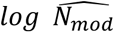 between cell types were tested with a *z*-test using the emmeans package, followed by Benjamini-Hochberg *P*-value correction for multiple testing (https://github.com/rvlenth/emmeans).

### Illumina RNA sequencing analysis

Reads processing was carried out using CGAT Flow pipelines [47]. Trim Galore was used for read trimming (https://github.com/FelixKrueger/TrimGalore). Trimmed reads were aligned using STAR [48], and quantified using featureCounts [49]. For snoRNA annotations, transcript annotations from hg38 human reference genome were combined with the snoRNA annotations from snoDB [8], using snoRupdate (https://github.com/scottgroup/snoRupdate).

Transcript expression was analysed using DESeq2 [50]. To map snoRNA to target rRNA positions, known snoRNA-rRNA interactions were extracted from snoDB [8].

## RESULTS

### Detecting rRNA modifications by Nanopore direct RNA sequencing

To characterise the rRNA modification profile during cell fate decisions, trilineage differentiation of hESCs into the three primary germ layers was used to model early embryonic gastrulation. hESC line MasterShef11 was differentiated into the three primary germ layers and RT-qPCR was performed to confirm the expression of pluripotency and germ layer marker genes (Fig S1). Ectoderm and mesoderm differentiation showed clear acquisition of new cell fates, while endoderm samples expressed both pluripotency and endodermal markers (Fig S1), as has previously been reported [12].

Libraries for RNA from pluripotent cells and each of the 3 lineages were prepared and sequenced using ONT RNA004 (Fig 1A). The reads obtained were demultiplexed and basecalled with the dorado package, using the super accurate mode which enabled basecalling of pseU and 2OMe modifications. The sequencing reads were then aligned to a human rRNA reference (Fig 1B). Each DRS library contained 90-209k reads for the 18S and 20-75k reads for the 28S (Table S2). Despite the initial RNA size selection, the low molecular weight 5.8S rRNA represented ∼50% of the resulting assigned reads, likely due to preferential sequencing of short RNAs by DRS.

**Figure 1:**
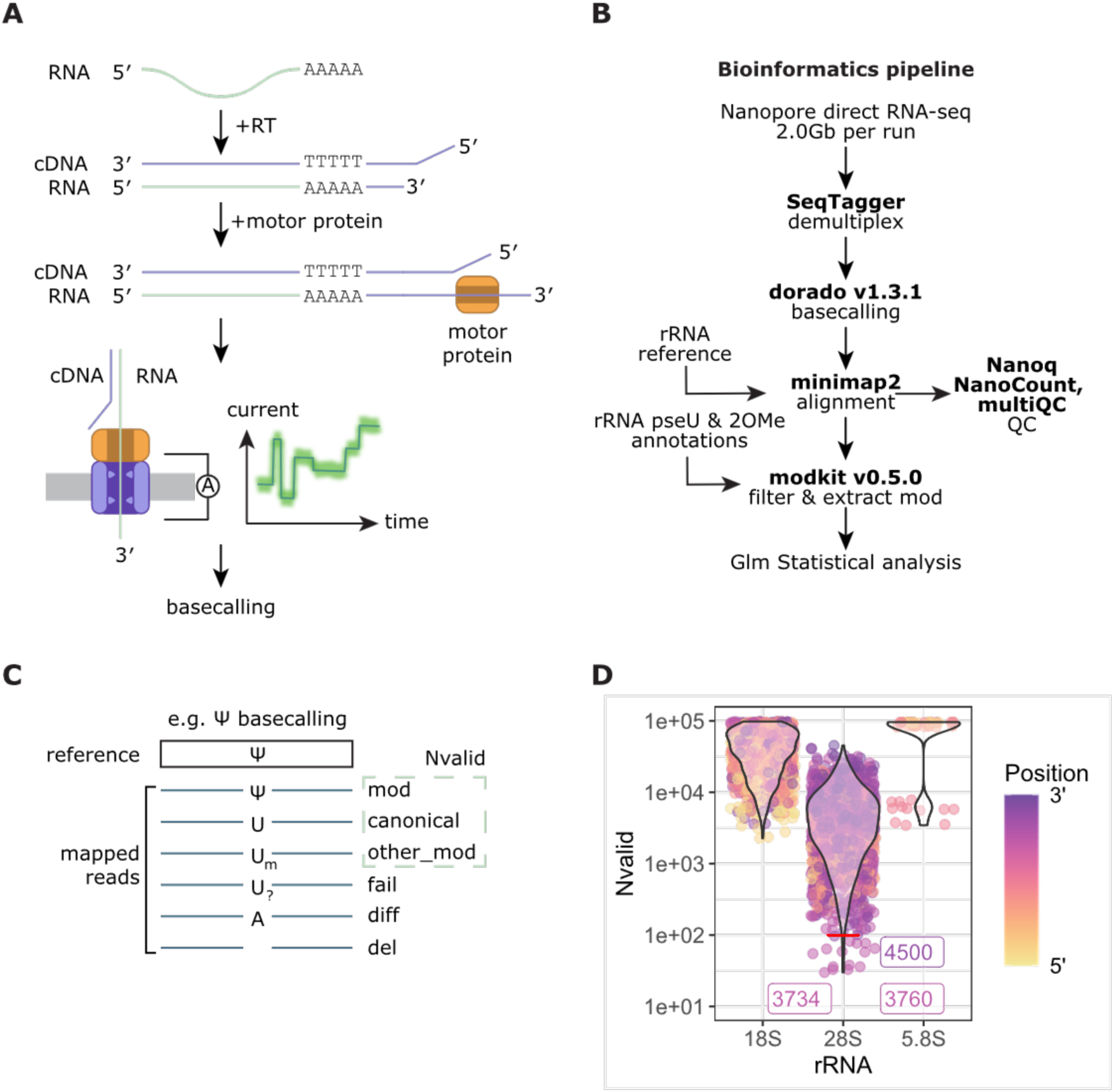
Overview of Nanopore direct RNA sequencing pipeline. **(A)** Principle of direct RNA sequencing. Total RNA was polyadenylated and ligated with an oligo(dT) adapter. Subsequent reverse transcription and ligation of sequencing motor protein allowed translocation of the RNA into a nanopore for sequencing. Electrical current across the nanopore represents a segment of RNA in the nanopore, which is then decoded and basecalled. **(B)** MShef11 total RNA were sequenced and mapped to a reference human rRNA sequence. Modification stoichiometry was calculated on all annotated pseU and 2OMe sites. Differential modification analysis between pluripotent and differentiated lineages was carried out using a generalised linear regression model (GLM). **(C)** Principle of basecall pileup. Basecalled nucleotides were filtered based on the corresponding probability score and assigned a modification type (see Figure S2). Nucleotides were then classified based on the assigned modification type and the annotated modification in the genome. The total number of nucleotides that matched with the reference base, and for which probability scores passed filtering, were defined as valid bases to quantify modification level. **(D)** Number of filtered valid bases per annotated modified position per sample. Sites are coloured by relative position along the length of the rRNA. Positions with Nvalid <100 are labelled.

Information of modified bases in the 18S, 28S, and 5.8S rRNA was extracted using the modkit package (Fig 1B), with each modified basecall being assigned a probability score. The modified basecalls were first processed by filtering basecalls with low probability. Using unmodified synthetic rRNAs as a negative control, probability thresholds were determined such that each rRNA species and each primary nucleotide has the same false positive rate in basecalling, which reduces the number of unmodified positions that were falsely assigned as modified (Fig S2A; Fig S2B; Table S3).

The number of filtered nucleotides at each modified position (Nvalid) was used to calculate modification levels (%mod = Nmod/Nvalid; Figure 1C). Assessing the Nvalid for each modified position found that sites on 18S and 5.8S rRNA both have Nvalid between 10,000-100,000, indicating that %mod calculated would be accurate. The 18S rRNA presented a modest 5′ bias where positions near the 5′ have a lower Nvalid (Fig 1D). This was likely due to DRS sequencing from the 3′ end, thus 3′ positions are preferentially sequenced and contain higher sequencing depth than 5′ positions. As the 28S rRNA had lower sequencing depth, the Nvalid across all modified positions on the 28S was lower compared to 18S and 5.8S (Fig 1D). Notably, there are three positions that have low Nvalid <100 bases. These positions did not appear to be 5′ or 3′ biased, but reside in proximity to other modified residues (Fig S3). Multiple modified positions in proximity may reduce modification detection accuracy and reduce modification probability, thus the Nvalid was also reduced.

### Ribosome heterogeneity through substoichiometric pseU and 2OMe modifications

Following filtering, the %mod of each modified position was calculated. To quantify %mod change during differentiation, a negative binomial regression was used to model the effects of cell types and rRNA position on modification stoichiometry. Pairwise z-tests were carried out between each germ layer and hESCs for each modified position (Fig S4). This approach effectively allows the fold change difference in %mod between two cell types to be assessed. As this statistical analysis was based on log ratios of Nmod, an absolute %mod change of 5% was implemented as a significance threshold to remove comparisons where both cell types exhibit low %mod, thus producing a high log ratio change, but a low absolute %mod change (Fig 2A). By comparing a single germ layer with hESC, differentiation-associated modification changes can be seen, with each germ layer presenting some significant modification changes (Fig 2A). Modified positions mainly demonstrated increased %mod, suggesting that reduced rRNA pseU and 2OMe may be a signature for pluripotency. The ectoderm and mesoderm also exhibited some positions with decreased %mod, but the endoderm did not.

**Figure 2:**
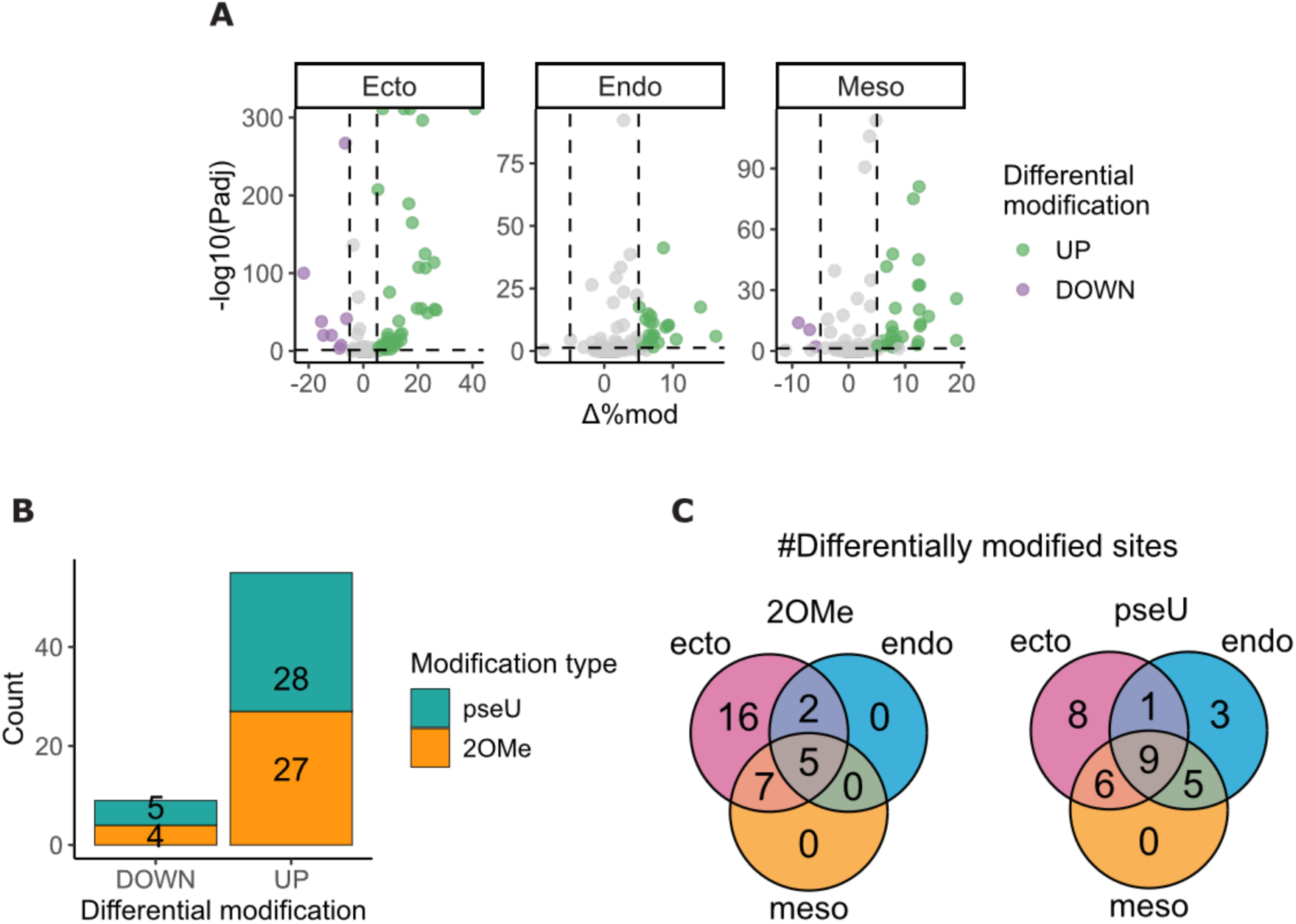
Differential modification analysis of 2′-*O*-methylated and pseudouridylated rRNA positions during differentiation. rRNA positions annotated to be modified by 2′-*O*-methylation and pseudouridylation were quantified by Nanopore direct RNA sequencing. Differential modification analysis was carried out on estimated log(Nmod) for each position between a differentiated cell type and hESCs, using a pairwise *z*-test. N = 3 biological replicates. **(A)** Volcano plot of differentially modified positions. Dashed lines indicate significance thresholds for absolute Δ%mod ≥ 5, and *z*-test adjusted *P* value < 0.05. Significantly differentially modified sites are coloured by direction of modification change from hESC. **(B)** The number of unique differentially modified positions categorised by the direction of modification change from hESC and the corresponding modification type. **(C)** The number of differentially modified positions distributed across the three primary germ layers.

pseU and 2OMe sites occupied a similar proportion of differentially modified sites, suggesting that modification type did not influence if a position was more or less likely to be differentially modified (Fig 2B). By comparing the distributions of differentially modified positions across the lineages, a large proportion of differentially modified sites were shared between multiple lineages (Fig 2C). While the ectoderm exhibited specific changes in 2OMe, the endoderm and mesoderm did not, as all differentially modified 2OMe sites were shared with the ectoderm. A similar pattern of differentially modified pseudouridylated sites was seen. While a few lineage specific modifications can be seen in the ectoderm and endoderm, the majority of differentially modified sites are found across multiple lineages. Differentially modified residues that were shared with multiple lineages suggest that the modification changes at these sites could be related to the exit of pluripotency, rather than a specific signature of differentiation to a defined lineage.

Statistical analysis of differential modification was visualised with %mod for all modified residues on each rRNA (Fig 3; Table S4). The modification levels of residues occupy the full range of %mod. Sites that were almost always modified (>85%) or rarely modified (<15%) show minimal variation between pluripotent and differentiated states. In contrast, positions with intermediate levels of modification (15-85%), representing 41% of pseU (43/104) and 35% of 2OMe (39/112), include numerous differentially modified sites (Fig 3). This broad distribution of fractional modification may indicate inherent modification heterogeneity across all rRNA species. Since a greater number of pseU sites, as compared to 2OMe, were fractionally modified, this may imply that pseU modifications may contribute more to intracellular ribosome heterogeneity.

**Figure 3:**
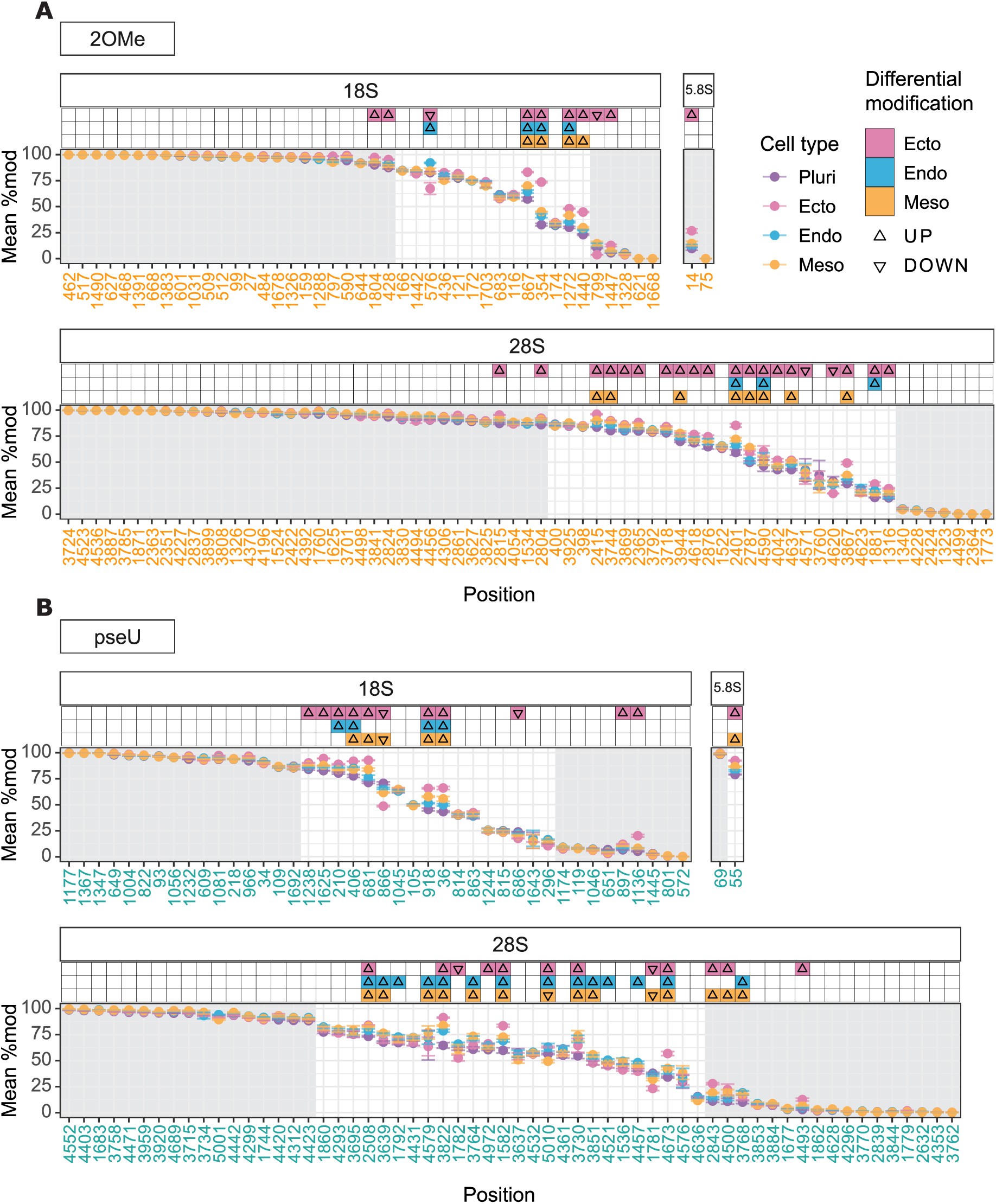
Modification levels of rRNA 2′-*O*-methylation and pseudouridylation in hESCs and trilineage differentiated cells. Annotated rRNA positions modified by **(A)** 2′-*O*-methylation and **(B)** pseudouridylation were quantified in the 18S, 5.8S, and 28S rRNA. Positions are arranged by decreasing %mod. Each point indicates the mean %mod ± 95% confidence interval for a single cell type. Positions with mean %mod < 15 or %mod > 85 in the pluripotent samples are shaded in grey. Tiles indicate pairwise differential modification analysis between a germ layer cell type and hESC. Coloured tiles indicate significant differential modification, defined as Δ%mod ≥ 5 and *z*-test of modification ratio where *P*_adj_ ≤ 0.05. N = 3 biological replicate.

The prevalence of differentially modified sites within this range of fractional modifications may indicate that heterogeneous ribosomes with substoichiometric modifications were also more likely to be regulated during differentiation. At positions where differential modification was observed, a common feature can be seen where %mod was lowest in hESCs, and showed a gradual increase from endoderm and mesoderm with the ectoderm having the highest %mod, where the increases ranged between 5-20% (Fig 3). A maximum of 40% increase was observed at 18S:Um354, similar to previous reports [29,30,31].

### Differential snoRNA expression during germ layer differentiation

While snoRNA expression levels do not generally correlate with levels of modification, previous studies have observed a positive correlation between dynamically modified 2OMe and snoRNA levels [10,11,12]. To determine if any of the differentially modified sites observed here also show a corresponding change in snoRNA expression during hESC differentiation, we next performed Illumina-based small RNA-seq.

Sequencing reads were mapped to snoDB annotations of human snoRNAs and differential expression analysis was carried out [8] (Table S5). snoRNAs that target rRNAs were then filtered for subsequent analysis. This detected 286 rRNA-modifying snoRNAs, which target 198/216 2OMe and pseU positions, as well as two modified positions of 1-methyl-3-α-amino-α- carboxyl-propylpseudouridine and 2′-*O*-methylpseudouridine (Fig S5). The remaining 18 2OMe and pseU positions have no snoRNA annotated in snoDB. Differential analysis of these snoRNAs, between differentiated cells and hESCs identified some to be differentially expressed (Fig 4A). Overall, the pattern of differential snoRNA expression was similar to that of rRNA modification with more snoRNAs showing an upregulation upon differentiation, as compared with those that are downregulated (Fig 4A). The majority of upregulated snoRNAs were C/D box snoRNAs, with H/ACA snoRNAs comprising one-third (Fig 4B). In contrast, the majority of down regulated snoRNA, were from the H/ACA class.

**Figure 4:**
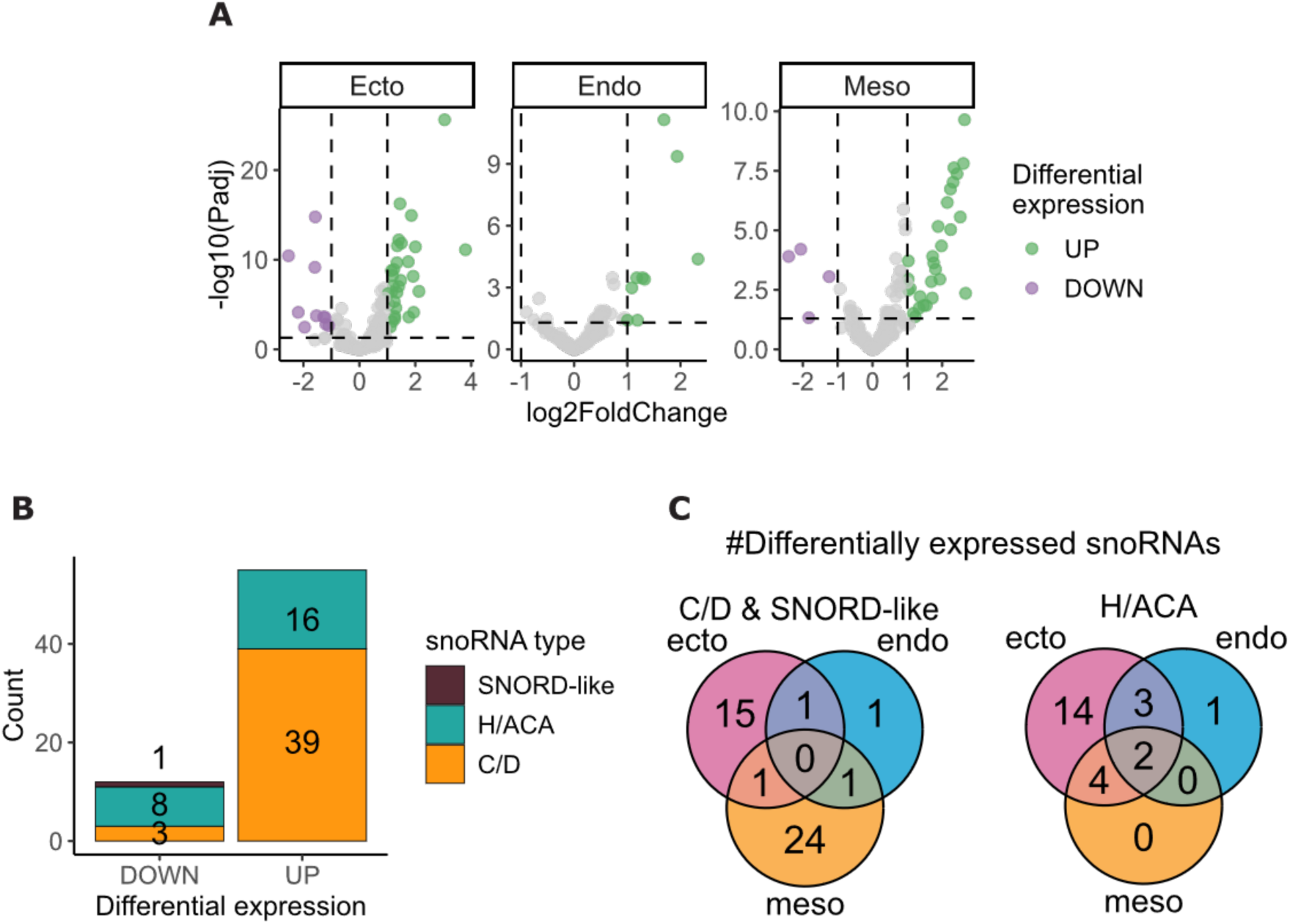
Differential snoRNA expression analysis during differentiation. snoRNA expressions were quantified by small RNA sequencing. Differential modification analysis was carried out on all mapped snoRNAs, and rRNA-targeting snoRNAs were selected. N = 4 biological replicates. **(A)** Volcano plot of differentially expressed snoRNAs. Each point indicates a pairwise comparison of a single snoRNA between a differentiated cell type and hESC. Dashed lines indicate significance thresholds for log_2_(fold change) in expression and Wald test adjusted *P* value. **(B)** The number of unique differentially expressed snoRNAs categorised by the direction of expression change from hESC and the corresponding snoRNA classes. **(C)** The number of differentially expressed snoRNAs distributed across the three primary germ layers.

The distribution of differentially expressed snoRNAs across the three lineages showed that C/D box snoRNAs were mainly lineage-specific, particularly for the ectoderm and mesoderm. Of the H/ACA box snoRNAs that were differentially expressed, most were associated specifically with the ectoderm with fewer found to be differentially expressed across multiple lineages (Fig 4C). This general pattern of lineage specific differential snoRNA expression contrasts with that seen for rRNA modification, which showed stoichiometric changes across multiple lineages.

### Some dynamic sites of rRNA modifications correlate with changes in guide snoRNA expressions

To determine if the differentially modified rRNA modification sites identified from DRS correlate with the differentially expressed snoRNAs, the datasets were integrated with respect to the snoRNA-rRNA interactions. Given that multiple snoRNAs may target a single rRNA position, the expressions of all snoRNAs that target the same rRNA site were summed, followed by differential log(fold change) of snoRNA expression to determine differential expression. Expression of snoRNAs that target multiple rRNA positions were assigned to each rRNA target analysed, such that rRNA modification levels were correlated with the maximum corresponding snoRNA expressions. Since this analysis relied on annotated snoRNA-rRNA interactions, rRNA sites with unknown snoRNA partners were excluded (Fig S5B).

Using the same significance thresholds for rRNA modification and snoRNA expression, nine sites demonstrated a correlation between snoRNA and rRNA expression. Seven sites were 2OMe, and two were pseU (Fig 5A). Consistent with the results from individual analyses of rRNA modification and snoRNA expression, ectoderm samples demonstrated the highest number of sites with correlated expressions. No sites were shared across multiple lineages (Fig 5A). Interestingly, the two pseU sites showed negative correlation between rRNA modification and guide snoRNA expression, as the fold change of summed snoRNA expression was negative while the change in %mod was positive (Fig 5A).

**Figure 5:**
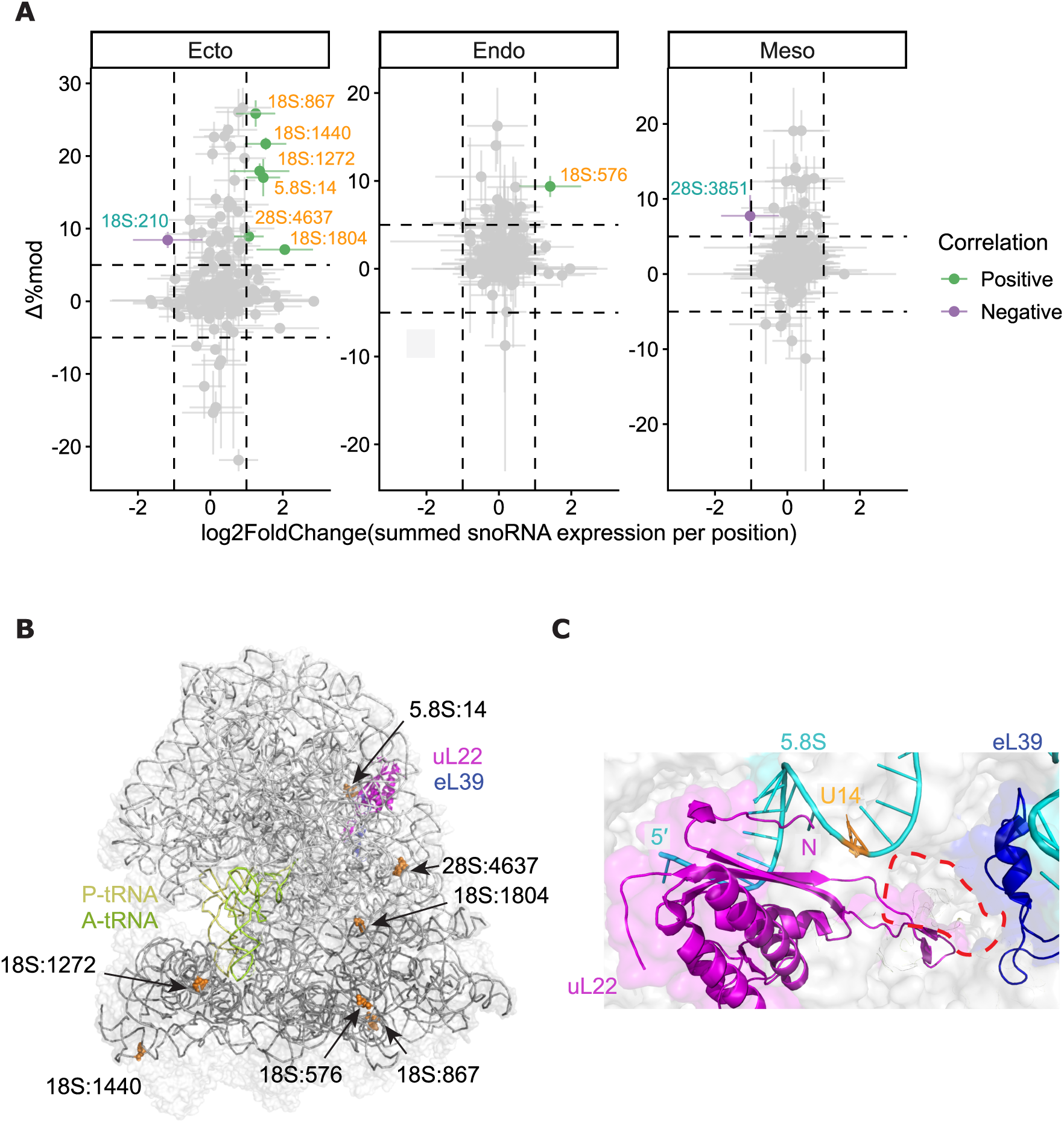
Integrated snoRNA expression-rRNA modification change during differentiation. **(A)** For each rRNA modified position, expression of all snoRNAs targeting the same position were summed, then log fold change was calculated relative to hESC. Dashed lines indicate significance thresholds for snoRNA expression and rRNA modification change. Error bars indicate ± 95% confidence interval. Points are coloured by correlation between rRNA %mod change and snoRNA log2 fold change. Sites with correlated response are labelled and coloured by modification type, 2OMe (orange) or pseU (teal). N = 3 or 4 biological replicates, for rRNA and snoRNA dataset respectively. **(B)** rRNA positions with positively correlated modification and guide snoRNA expression highlighted as spheres on the structure of a human ribosome (PDB:8JDK). rRNA and tRNA are shown as grey and yellow/green ribbons respectively, and the peptide exit channel proteins uL22 and eL39 are highlighted in magenta and blue respectively. **(C)** Magnified view of the peptide exit channel (red dashed circle). The 5.8S rRNA is highlighted in cyan, with position U14 shown in orange. The N-terminus of uL22 and the 5′ end of the 5.8S rRNA are labelled, and r-proteins are coloured as in **B**.

### Dynamic modifications with correlated snoRNA expression may indicate sites of translational regulation

The seven 2OMe sites with increased %mod and correlated guide snoRNA expressions consist of five positions in the ribosome small subunit, and two positions in the large subunit. The location of these sites may indicate potential regulatory capacity from the presence/absence of the modification through proximity to functionally important regions of the ribosome (Fig 5B). One example, 18S:1804, maps to the subunit interface, suggesting that its modification may influence subunit joining or ribosome translocation. Similarly, 5.8S:14 lies near the peptide exit channel proteins eL39 and the N-terminus of uL22, where the presence or absence of modification may impact how uL22 interacts with the nascent peptide during translation elongation (Fig 5C). The localisation of these differentially modified nucleotides at key positions on the ribosome, highlights them as potential targets for specialised translational regulation during cell fate decisions.

## DISCUSSION

Here we describe the use of Nanopore DRS to comprehensively analyse the pseU and 2OMe modification landscape of rRNA in hESCs and their differentiated lineages. Modification profiling revealed that approximately 59-65% of annotated pseU and 2OMe sites were either heavily (>85%) or rarely (<15%) modified, and exhibited minimal variation during differentiation. In contrast, among the remaining sites displaying intermediate levels of modification (15-85%), we identified an enrichment of residues that undergo significant changes in modification stoichiometry upon differentiation, with a high degree of overlap between lineages. These findings highlight that the substoichiometrically modified sites represent the most dynamically regulated positions within the rRNA, and potentially mediate an exit programme from pluripotency.

These analyses demonstrate distinct patterns of individual base modifications that give rise to rRNA heterogeneity. While a subset of these sites show differential modification during differentiation and support the concept of intercellular ribosome heterogeneity, a large proportion of the substoichiometrically modified sites did not, indicating the presence of intracellular rRNA subtypes [11,40]. This basal pool of heterogeneous ribosomes may represent a complex ensemble of distinct ribosome subpopulations that may facilitate specialised translational programmes. Interestingly, while over 40% (54/134) of the heavily modified sites are conserved across eukaryotes (when compared to *S. cerevisiae*), only 22% (18/82) of sites exhibiting intermediate levels of modification are conserved (Table S4). This suggests that greater flexibility in modification status, and thus intra/intercellular heterogeneity, resides in non-conserved sites.

Previous quantitative sequencing studies investigating pseU and 2OMe modifications have relied heavily on Illumina-based methods such as RiboMeth-seq and HydraPsi-seq for detection [34,35]. These approaches typically depend on chemical treatments that induce read truncation or specific base conversion at the modified residue, to infer modification status through cDNA sequencing [36]. However, with recent advances in DRS, pseU and 2OMe can be simultaneously detected and basecalled directly, effectively circumventing the potential limitations and biases of indirect methods. Furthermore, the use of an unmodified synthetic rRNA control allows modification calling to be assessed in a context-specific manner. By defining an empirical false positive threshold within the control samples, filtering criteria for modified basecalls can be tailored to the specific sequence context of the rRNA, ensuring high confidence basecalls are retained in the biological datasets [51]. In addition, our statistical analysis of differential modification relies on modelling read count distributions, similar to common RNA-seq differential expression analysis, thereby providing a more accurate assessment of stoichiometric changes between conditions.

A comparison of the DRS derived patterns of 2OMe reported here with earlier findings from RiboMeth-seq studies of human rRNA shows that 2OMe levels identified in RiboMeth-seq are generally higher than that seen in DRS analyses [9,11,12,30]. Further, RiboMeth-seq shows less site-to-site variation in levels of 2OMe, categorising approximately two-thirds of all 2OMe sites highly modified [9,12]. Despite the differences in the magnitude of modification scores between DRS and RiboMeth-seq, similar conclusions could be drawn from differential analyses. One RiboMeth-seq study investigated 2OMe modification heterogeneity during trilineage differentiation of H9 hESCs, using a similar differentiation strategy to ours, with the exception of the ectoderm, which was progressively differentiated to early/late neural progenitors and mature neurons [12]. Out of the 22 differentially modified 2OMe sites reported from RiboMeth-seq, 14 were also captured in our DRS dataset [12] (Figure S6). Since most of the differentially modified sites we have detected arise in the ectoderm, the remaining sites (8 sites unique to RiboMeth-seq and 16 sites unique to this study) likely reflect the distinct temporal stages of ectoderm differentiation analysed. This variation suggests that the modification signature may change at different timepoints within a lineage and highlights the effectiveness of DRS in capturing highly dynamic rRNA modification states.

Unlike 2OMe, pseU heterogeneity has not previously been explored for hESC fate decisions. While HydraPsi-seq has identified heterogeneity in multiple human cell lines, as well as in SH- SY5Y neuroblastoma differentiation, no quantitative analyses were performed to compare proliferative and differentiated states [10,35]. In addition to Illumina-based sequencing methods, earlier versions of DRS (RNA002) that lacked modification basecalling leveraged on error rates to infer modification status. rRNA modifications identified in this manner demonstrated modification heterogeneity during neuronal differentiation of mESCs, highlighting the presence of dynamic rRNA modification patterns across cell fates [31]. In more recent versions of DRS (dorado v0.7), basecalling of rRNA modifications became available, beginning with pseU, which has enabled the analysis of tracking pseU modifications during rRNA processing in HEK293T cells [52]. The patterns of pseU modifications in our MShef11 dataset (using dorado v1.3) largely resembled what was detected in HEK293T, reinforcing the observation of pseU heterogeneity in rRNA [52]. Overall, the results presented here are consistent with other studies exploring modification patterns using earlier versions of DRS.

While previous studies have identified multiple sites where levels of modification fluctuate during cell-fate decisions, few have been directly shown to play a functional role [11,12]. Characterized examples of 2OMe sites with specialised regulatory functions exhibited two characteristics: a large magnitude of %mod change among all profiled sites, and a positive correlation between %mod and guide snoRNA expression [11,12]. Integration of our DRS with snoRNA-seq revealed little global correlation between datasets, as has previously been described [9,10]. However, seven 2OMe positions showed positive correlation between differential snoRNA expression and rRNA modification changes during differentiation. Notably, out of the seven differentially modified 2OMe sites identified in Figure 5A, five (excluding 18S:1804 and 28S:4637) overlap with sites known to undergo cell fate associated modification change [12] (Figure S6). These consistent findings further support that the candidate sites identified through integrating DRS and snoRNA-seq could mediate specialised translation.

Among the correlated sites, the modification 5.8S:U14 is particularly noteworthy. This site is located proximal to the peptide exit channel, a region previously shown to exhibit structural heterogeneity in pluripotent cells, through the incorporation of the mammalian-specific r-protein paralogue eL39L [19,53] (Figure 5C). eL39L is upregulated in pluripotent and male germ cells, where it modulates co-translational protein folding [19,53]. Given the proximity of 5.8S:U14 and eL39/L, combined with the known heterogeneity at the 5’ end of 5.8S rRNA (long and short forms) [54,55], it is tempting to speculate that heterogeneity in this region may be co-regulated. Such coordinated variation may serve to fine tune the functional dynamics of the peptide exit channel.

Overall, our findings provide a high-resolution map of the human rRNA landscape during hESC differentiation and further strengthens the growing evidence of rRNA modification heterogeneity both within cells and between cell types. Through the direct profiling of modifications via DRS, we show that sites of substoichiometric modification serve as key areas of dynamic regulation during cell fate decisions. Further, the analysis pipeline developed provides a standard for quantifying modification levels with high stringency, which will facilitate analyses of the DRS datasets and future investigations into modification heterogeneity.

## ACKNOWLEDGEMENTS

We thank Anders Lund (University of Copenhagen) for the kind gift of the pcDNA3.1 plasmids containing human 5S, 5.8S, 18S, and 28S rRNA. We thank Owen Laing and Gabriele Gelezauskaite in the Barbaric lab for assistance in low pass sequencing. We acknowledge IT Services at The University of Sheffield for the provision of High Performance Computing services.

## AUTHOR CONTRIBUTIONS

Tessa Chan: Conceptualisation, Data curation, Formal analysis, Investigation, Methodology, Validation, Visualization, Writing – original draft, Writing – revision and editing. Ivana Barbaric: Resources, Supervision, Writing – reviewing and editing. Emma Thomson: Conceptualisation, Formal analysis, Funding acquisition, Project administration, Resources, Supervision, Writing – original draft, Writing – revision and editing.

## SUPPLEMENTARY DATA

Supplementary Data are available at NAR online.

## DATA AVAILABILITY

Raw FASTQ files from small RNA-sequencing, and basecalled BAM files from DRS generated in this study are available in GEO under accessions GSE335337 and GSE336742, respectively. The list of annotated rRNA modified residues was collated from snoDB [8] and Taoka *et al.* (2018) [3]. Annotations of conserved rRNA pseU/2OMe residues between human and yeast was taken from snoRNABase [56]. The reference sequences of rRNA, modified residue annotations, analysis scripts, conda environment YML files, and apptainer DEF files are available in this project’s Github repository (https://github.com/Thomson-RNA-Lab/Integrated-rRNA-snoRNA-seq).

## REFERENCES

1. Decatur, W.A. and Fournier, M.J. (2002) rRNA modifications and ribosome function. Trends Biochem. Sci., 27:344–351. doi: 10.1016/s0968-0004(02)02109-6.

2. Natchiar, S.K., Myasnikov, A.G., Kratzat, H. et al. (2017) Visualization of chemical modifications in the human 80S ribosome structure. Nature, 551:472–477. doi: 10.1038/nature24482.

3. Taoka, M., Nobe, Y., Yamaki, Y. et al. (2018) Landscape of the complete RNA chemical modifications in the human 80S ribosome. Nucleic Acids Res., 46:9289–9298. doi: 10.1093/nar/gky811.

4. Balakin, A.G., Smith, L. and Fournier, M.J. (1996) The RNA world of the nucleolus: two major families of small RNAs defined by different box elements with related functions. Cell, 86:823– 834. doi: 10.1016/s0092-8674(00)80156-7.

5. Kiss-László, Z., Henry, Y., Bachellerie, J.P. et al. (1996) Site-specific ribose methylation of preribosomal RNA: a novel function for small nucleolar RNAs. Cell, 85:1077–1088. doi: 10.1016/s0092-8674(00)81308-2.

6. Bortolin, M.L., Ganot, P. and Kiss, T. (1999) Elements essential for accumulation and function of small nucleolar RNAs directing site-specific pseudouridylation of ribosomal RNAs. EMBO J., 18:457–469. doi: 10.1093/emboj/18.2.457.

7. Jorjani, H., Kehr, S., Jedlinski, D.J. et al. (2016) An updated human snoRNAome. Nucleic Acids Res., 44:5068–5082. doi: 10.1093/nar/gkw386.

8. Bergeron, D., Paraqindes, H., Fafard-Couture, É. et al. (2022) snoDB 2.0: an enhanced interactive database, specializing in human snoRNAs. Nucleic Acids Res., 51:D291–D296. doi: 10.1093/nar/gkac835.

9. Krogh, N., Jansson, M.D., Häfner, S.J. et al. (2016) Profiling of 2ʹ-O-Me in human rRNA reveals a subset of fractionally modified positions and provides evidence for ribosome heterogeneity. Nucleic Acids Res., 44:7884–7895. doi: 10.1093/nar/gkw482.

10. Gawade, K., Plewka, P., Häfner, S.J. et al. (2023) FUS regulates a subset of snoRNA expression and modulates the level of rRNA modifications. Sci. Rep., 13:2974. doi: 10.1038/s41598-023-30068-2.

11. Jansson, M.D., Häfner, S.J., Altinel, K. et al. (2021) Regulation of translation by site-specific ribosomal RNA methylation. Nat. Struct. Mol. Biol., 28:889–899. doi: 10.1038/s41594-021-00669-4.

12. Häfner, S.J., Jansson, M.D., Altinel, K. et al. (2023) Ribosomal RNA 2ʹ-O-methylation dynamics impact cell fate decisions. Dev. Cell, 58:1593–1609.e9. doi: 10.1016/j.devcel.2023.06.007.

13. Ingolia, N., Lareau, L. and Weissman, J. (2011) Ribosome profiling of mouse embryonic stem cells reveals the complexity and dynamics of mammalian proteomes. Cell, 147:789–802. doi: 10.1016/j.cell.2011.10.002.

14. Friend, K., Brooks, H.A., Propson, N.E. et al. (2015) Embryonic stem cell growth factors regulate eIF2α phosphorylation. PLoS One, 10:e0139076. doi: 10.1371/journal.pone.0139076.

15. Gabut, M., Bourdelais, F. and Durand, S. (2020) Ribosome and translational control in stem cells. Cells, 9:497. doi: 10.3390/cells9020497.

16. Guo, H. (2018) Specialized ribosomes and the control of translation. Biochem. Soc. T., 46:855– 869. doi: 10.1042/bst20160426.

17. Beavan, A.J.S., Thuburn, V., Fatkhullin, B. et al. (2025) Specialized ribosomes: integrating new insights and current challenges. Philos. Trans. R. Soc. B, 380:20230377. doi: 10.1098/rstb.2023.0377.

18. Hopes, T., Norris, K., Agapiou, M. et al. (2022) Ribosome heterogeneity in Drosophila melanogaster gonads through paralog-switching. Nucleic Acids Res., 50(4):2240–2257. doi: 10.1093/nar/gkab606.

19. Li, H., Huo, Y., He, X. et al. (2022) A male germ-cell-specific ribosome controls male fertility. Nature, 612(7941):725–731. doi: 10.1038/s41586-022-05508-0.

20. Shiraishi, C., Matsumoto, A., Ichihara, K. et al. (2023) RPL3L-containing ribosomes determine translation elongation dynamics required for cardiac function. Nat. Commun., 14:2131. doi: 10.1038/s41467-023-37838-6.

21. Shi, Z., Fujii, K., Kovary, K.M. et al. (2017) Heterogeneous ribosomes preferentially translate distinct subpools of mRNAs genome-wide. Mol. Cell, 67:71–83.e7. doi: 10.1016/j.molcel.2017.05.021.

22. Genuth, N.R., Shi, Z., Kunimoto, K. et al. (2022) A stem cell roadmap of ribosome heterogeneity reveals a function for RPL10A in mesoderm production. Nat. Commun., 13(1):5491. doi: 10.1038/s41467-022-33263-3.

23. Dopler, A., Alkan, F., Malka, Y. et al. (2024) P-stalk ribosomes act as master regulators of cytokine-mediated processes. Cell, 187:6981–6993.e23. doi: 10.1016/j.cell.2024.09.039.

24. Imami, K., Milek, M., Bogdanow, B. et al. (2018) Phosphorylation of the ribosomal protein RPL12/uL11 affects translation during mitosis. Mol. Cell, 72:84–98.e9. doi: 10.1016/j.molcel.2018.08.019.

25. Locati, M.D., Pagano, J.F.B., Girard, G. et al. (2017) Expression of distinct maternal and somatic 5.8S, 18S, and 28S rRNA types during zebrafish development. RNA, **23**:1188–1199. doi: 10.1261/rna.061515.117.

26. Parks, M.M., Kurylo, C.M., Dass, R.A. et al. (2018) Variant ribosomal RNA alleles are conserved and exhibit tissue-specific expression. Sci. Adv., 4:eaao0665. doi: 10.1126/sciadv.aao0665.

27. Rothschild, D., Susanto, T.T., Sui, X. et al. (2024) Diversity of ribosomes at the level of rRNA variation associated with human health and disease. Cell Genomics, 4:100629. doi: 10.1016/j.xgen.2024.100629.

28. Moser, T.V., Bond, D.M. and Hore, T.A. (2025) Variant ribosomal DNA is essential for female differentiation in zebrafish. Philos. Trans. R. Soc. B, 380:20240107. doi: 10.1098/rstb.2024.0107.

29. Hebras, J., Krogh, N., Marty, V. et al. (2019) Developmental changes of rRNA ribose methylations in the mouse. RNA Biol., 17:150–164. doi: 10.1080/15476286.2019.1670598.

30. Krogh, N., Asmar, F., Côme, C. et al. (2020) Profiling of ribose methylations in ribosomal RNA from diffuse large B-cell lymphoma patients for evaluation of ribosomes as drug targets. NAR Cancer, 2:zcaa035. doi: 10.1093/narcan/zcaa035.

31. Milenkovic, I., Cruciani, S., Llovera, L. et al. (2025) Epitranscriptomic rRNA fingerprinting reveals tissue-of-origin and tumor-specific signatures. Mol. Cell, 85:177–190.e7. doi: 10.1016/j.molcel.2024.11.014.

32. Carlile, T.M., Rojas-Duran, M.F., Zinshteyn, B. et al. (2014) Pseudouridine profiling reveals regulated mRNA pseudouridylation in yeast and human cells. Nature, 515:143–146. doi: 10.1038/nature13802.

33. Schwartz, S., Bernstein, D.A., Mumbach, M.R. et al. (2014) Transcriptome-wide Mapping Reveals Widespread Dynamic-Regulated Pseudouridylation of ncRNA and mRNA. Cell, 159:148–162. doi: 10.1016/j.cell.2014.08.028.

34. Birkedal, U., Christensen-Dalsgaard, M., Krogh, N. et al. (2014) Profiling of ribose methylations in RNA by high-throughput sequencing. Angew. Chem. Int. Ed., 54:451–455. doi: 10.1002/anie.201408362.

35. Marchand, V., Pichot, F., Neybecker, P. et al. (2020) HydraPsiSeq: a method for systematic and quantitative mapping of pseudouridines in RNA. Nucleic Acids Res., 48:e110–e110. doi: 10.1093/nar/gkaa769.

36. Zhang, Y., Lu, L. and Li, X. (2022) Detection technologies for RNA modifications. Exp. Mol. Med., 54:1601–1616. doi: 10.1038/s12276-022-00821-0.

37. Liu, H., Begik, O., Lucas, M.C. et al. (2019) Accurate detection of m6A RNA modifications in native RNA sequences. Nat. Commun., 10:4079. doi: 10.1038/s41467-019-11713-9.

38. Begik, O., Lucas, M.C., Pryszcz, L.P. et al. (2021) Quantitative profiling of pseudouridylation dynamics in native RNAs with nanopore sequencing. Nat. Biotechnol., 39:1278–1291. doi: 10.1038/s41587-021-00915-6.

39. Leger, A., Amaral, P.P., Pandolfini, L. et al. (2021) RNA modifications detection by comparative Nanopore direct RNA sequencing. Nat. Commun., 12:7198. doi: 10.1038/s41467-021-27393-3.

40. Bailey, A.D., Talkish, J., Ding, H. et al. (2022) Concerted modification of nucleotides at functional centers of the ribosome revealed by single-molecule RNA modification profiling. eLife, 11:e76562. doi: 10.7554/elife.76562.

41. Acera Mateos, P., J Sethi, A., Ravindran, A., et al. (2024) Prediction of m6A and m5C at single-molecule resolution reveals a transcriptome-wide co-occurrence of rna modifications. Nat. Commun., 15:3899. doi: 10.1038/s41467-024-47953-7.

42. Pryszcz, L.P., Diensthuber, G., Llovera, L. et al. (2025) Rapid and accurate demultiplexing of direct RNA nanopore sequencing data with SeqTagger. Genome Res., 35:956–966. doi: 10.1101/gr.279290.124.

43. Li, H. (2018) Minimap2: pairwise alignment for nucleotide sequences. Method. Biochem. Anal., 34:3094–3100. doi: 10.1093/bioinformatics/bty191.

44. Gleeson, J., Leger, A., Prawer, Y.D.J. et al. (2021) Accurate expression quantification from nanopore direct RNA sequencing with NanoCount. Nucleic Acids Res., 50:e19. doi: 10.1093/nar/gkab1129.

45. Steinig, E. and Coin, L. (2022) Nanoq: ultra-fast quality control for nanopore reads. J. Open Source Softw., 7:2991. doi: 10.21105/joss.02991.

46. Ewels, P., Magnusson, M., Lundin, S. et al. (2016) MultiQC: summarize analysis results for multiple tools and samples in a single report. Method. Biochem. Anal., 32:3047–3048. doi: 10.1093/bioinformatics/btw354.

47. Cribbs, A.P., Luna-Valero, S., George, C. et al. (2019) CGAT-core: a python framework for building scalable, reproducible computational biology workflows. F1000Research, **8**:377. doi: 10.12688/f1000research.18674.1.

48. Dobin, A., Davis, C.A., Schlesinger, F. et al. (2012) STAR: ultrafast universal RNA-seq aligner. Bioinformatics, 29:15–21. doi: 10.1093/bioinformatics/bts635.

49. Liao, Y., Smyth, G.K. and Shi, W. (2013) featureCounts: an efficient general purpose program for assigning sequence reads to genomic features. Bioinformatics, 30:923–930. doi: 10.1093/bioinformatics/btt656.

50. Love, M.I., Huber, W. and Anders, S. (2014) Moderated estimation of fold change and dispersion for RNA-seq data with DESeq2. Genome Biol., 15:550. doi: 10.1186/s13059-014-0550-8.

51. Cruciani, S. and Novoa, E.M. (2025) The new era of single-molecule RNA modification detection through nanopore base-calling models. Nat. Rev. Mol. Cell Bio., 27:10–18. doi: 10.1038/s41580-025-00896-3.

52. Pastore, S., Wacheul, L., Lehmann, L. et al. (2026) Mapping human pre-rRNA processing and modification at single nucleotide resolution using long read nanopore sequencing. Nat. Commun., 17:4658. doi: 10.1038/s41467-026-71164-x.

53. Banerjee, A., Ataman, M., Smialek, M.J. et al. (2024) Ribosomal protein RPL39L is an efficiency factor in the cotranslational folding of a subset of proteins with alpha helical domains. Nucleic Acids Res., 52(15):9028–9048. doi: 10.1093/nar/gkae630.

54. Wang, M., Parshin, A.V., Shcherbik, N. et al. (2015) Reduced expression of the mouse ribosomal protein Rpl17 alters the diversity of mature ribosomes by enhancing production of shortened 5.8S rRNA. RNA, 21:1240–1248. doi: 10.1261/rna.051169.115.

55. Venturi, G., Zacchini, F., Ruzzi, F. et al. (2025) 5.8S rRNA forms and ribosome heterogeneity in breast cancer. Biochimie, 238:57–64. doi: 10.1016/j.biochi.2025.05.010.

56. Lestrade, L. and Weber, M.J. (2006) snoRNA-LBME-db, a comprehensive database of human H/ACA and C/D box snoRNAs. Nucleic Acids Res., 34:D158–D162. doi: 10.1093/nar/gkj002.

