## Supplemental data for "Integrated sequencing approach to probe rRNA modification landscape during human embryonic stem cell differentiation"

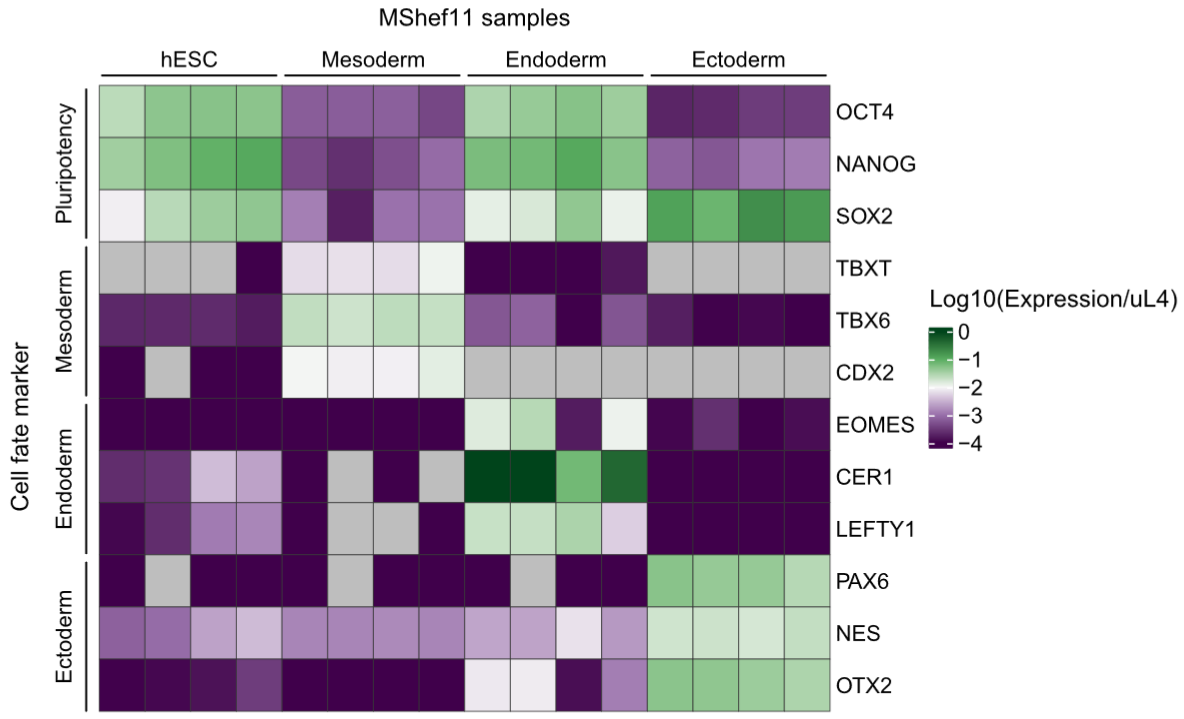


**Figure S1:** mRNA expression of known cell fate marker genes, detected by RT-qPCR. Each column represents a biological replicate of a single cell type. RNA expression was normalised to ribosomal protein *uL4* as a housekeeping gene, N = 4 biological replicates.

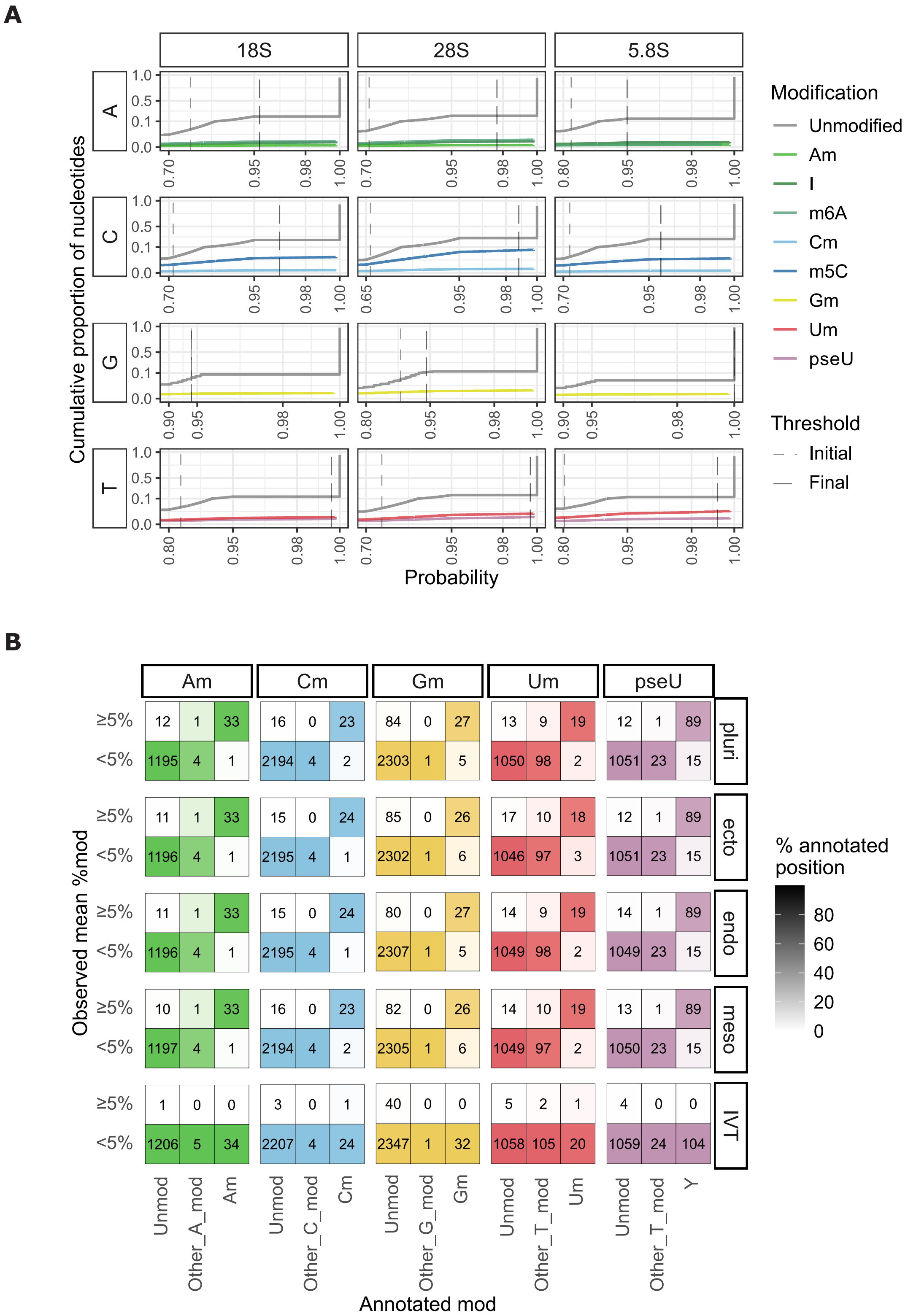


**Figure S2:** Probability threshold for filtering modified basecalls. (A) Probability distribution of unfiltered basecalls in the IVT samples. Separate thresholds were set for each canonical base and rRNA. Vertical lines indicate the default threshold (light grey) and the final threshold (dark grey) used. (B) The number of positions categorised by the corresponding annotated modification and the observed mean modification levels. Each vertical facet indicates the basecalling model, and each horizontal facet indicates the sample type. Within each facet of sample type and basecalling model, each cell is shaded according to the proportion of annotated positions in each observed %mod category. N = 2 or 3 for IVT and MShef11 samples respectively.

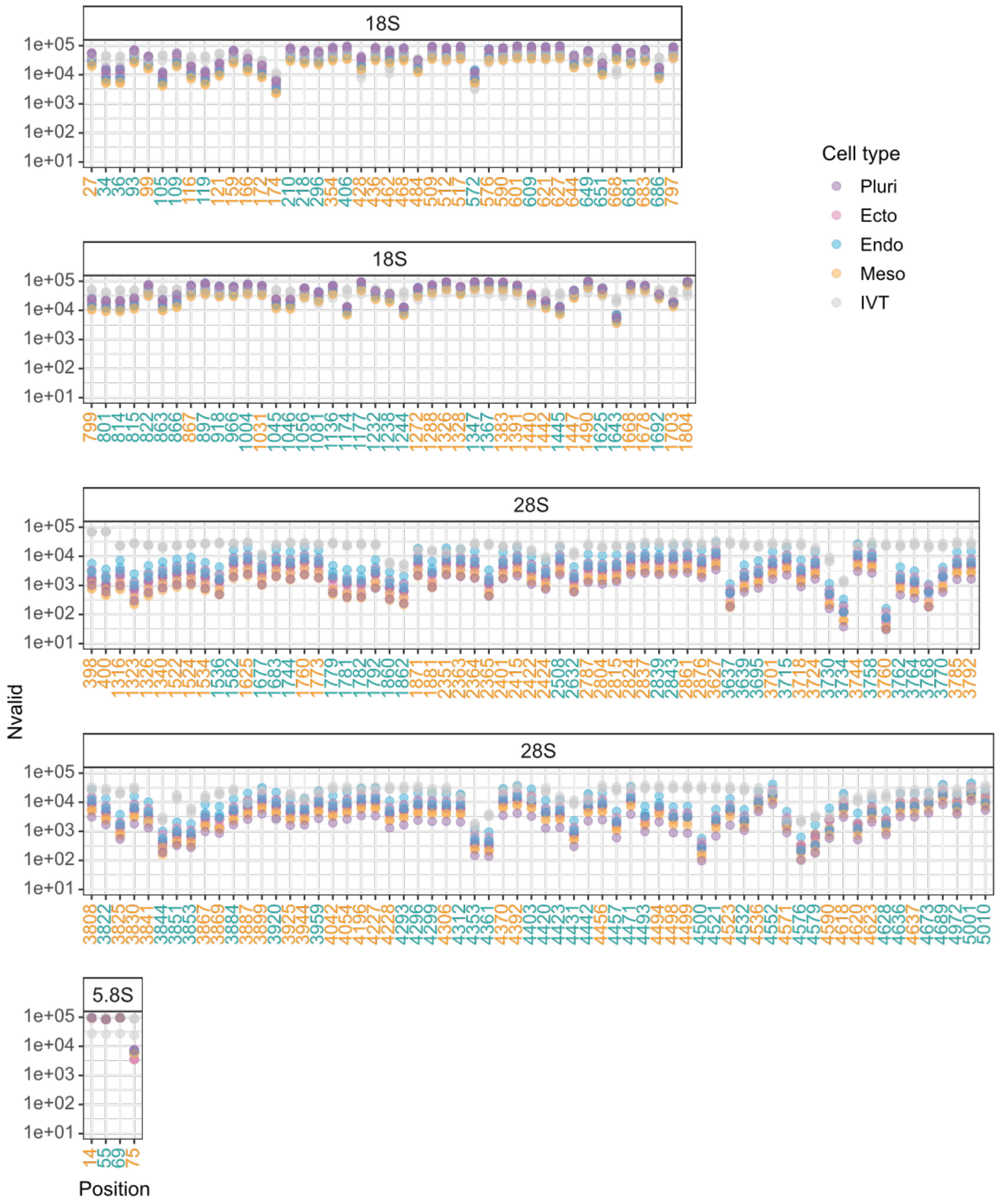


**Figure S3:** Number of filtered basecalls (Nvalid) used to quantify modification stoichiometry by DRS. All known pseU (cyan) and 2OMe (orange) positions in the rRNA are plotted from 5′–3′. N = 2 or 3 for IVT and MShef11 samples respectively.

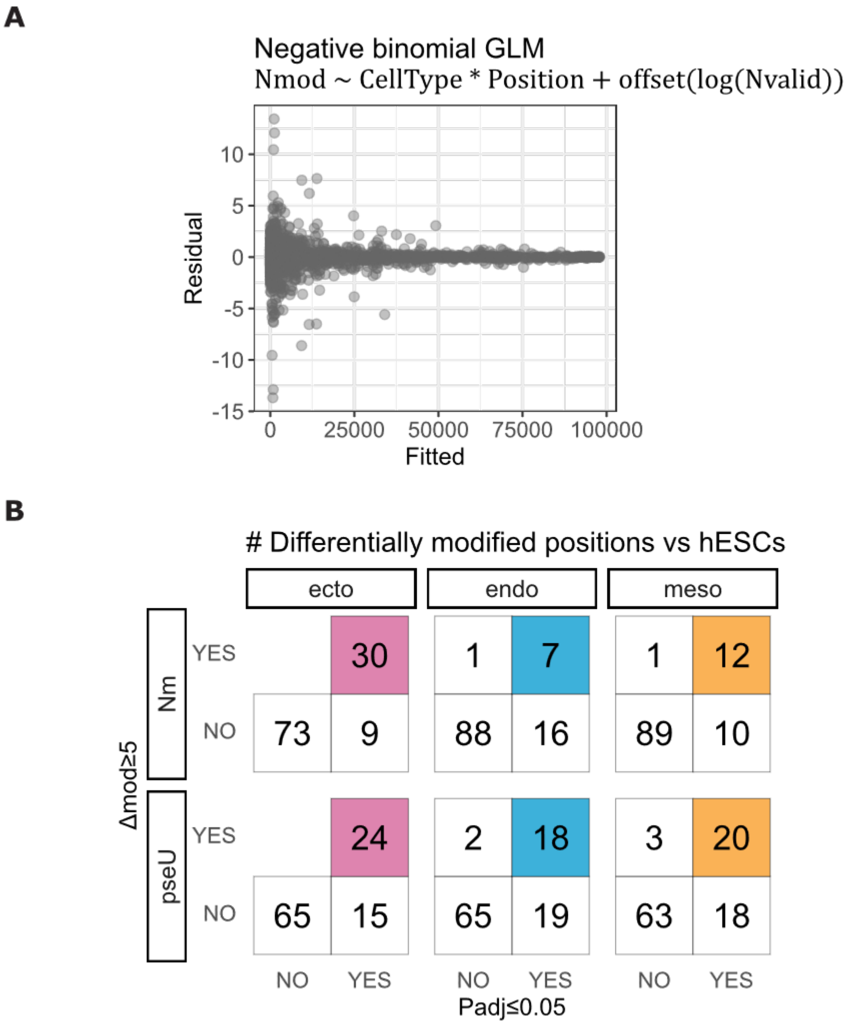


**Figure S4:** Statistical modelling of differential modification for MShef11 DRS. (A) Residual plot of the negative binomial GLM used to model the number of modified basecalls (Nmod) as a function of cell type and position, offset by the total number of valid basecalls (Nvalid). (B) The number of differentially modified positions filtered by statistical testing and absolute modification changes, for each pairwise comparison between individual MShef11 differentiated cell types and hESCs. N = 3 biological replicates.

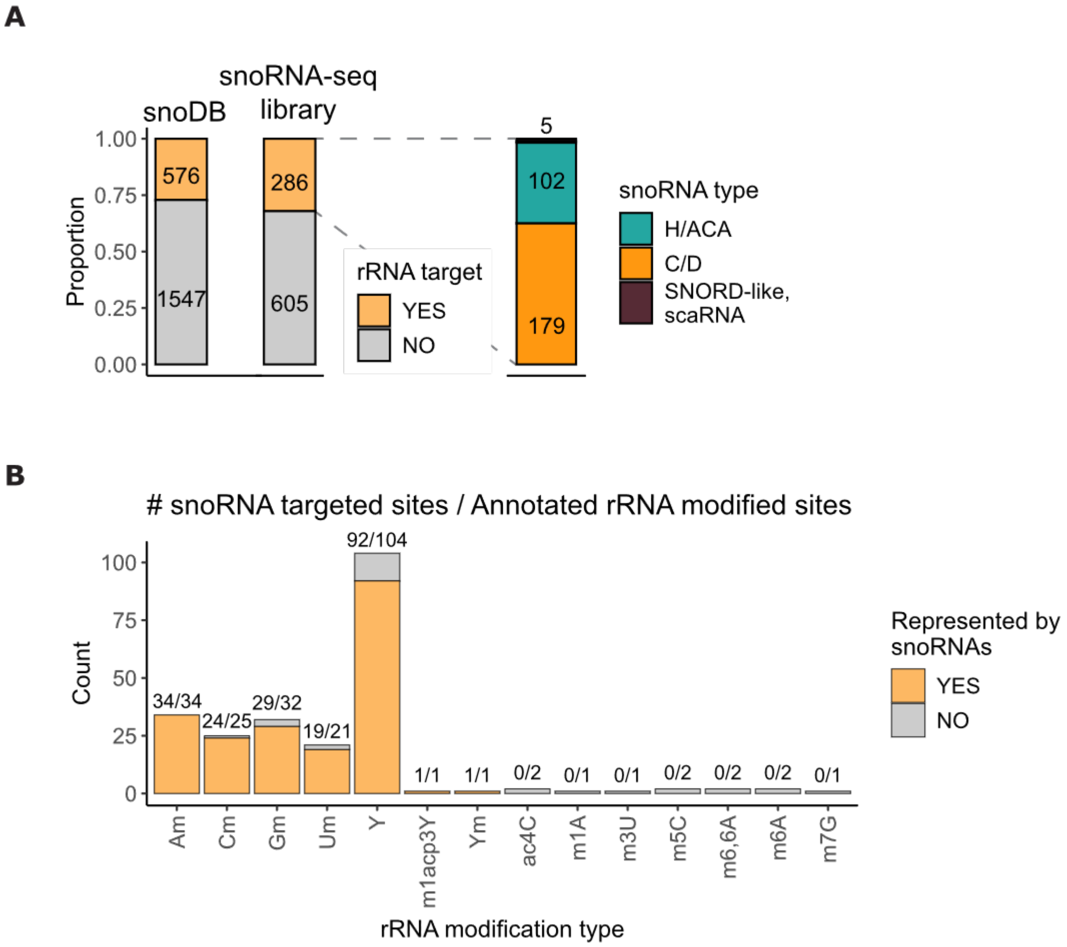


**Figure S5:** Summary of rRNA-targeting snoRNAs from MShef11 snoRNA-seq. (A) The number of snoRNA genes mapped by snoRNA-seq was first categorised by whether the snoRNA targets rRNA (left). The rRNA-targeting snoRNAs were further classified by the snoRNA types (right). (B) The number of unique rRNA modified sites targeted by snoRNAs from the snoRNA-seq dataset. N = 4 biological replicates.

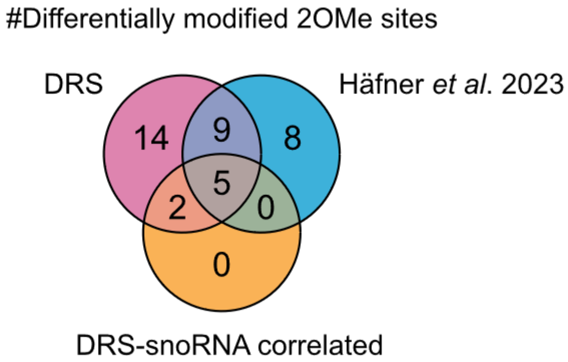


**Figure S6:** The number of differentially modified 2OMe sites in the MShef11 DRS dataset and in the H9 RiboMeth-seq dataset, across all lineage differentiation.

**Table S1:** Oligonucleotides used in this study

| Name | 5′–3′ sequence | Usage |
|---|---|---|
| HsGAPDH_qPCR_Fw1 | CTTTTGCGTCGCCAGCC | RT-qPCR |
| HsGAPDH_qPCR_Rv1 | GACCAGGCGCCCAATACG | RT-qPCR |
| HsOCT4_qPCR_Fw3 | CCCACACTGCAGCAGATCA | RT-qPCR |
| HsOCT4_qPCR_Rv3 | ACCACACTCGGACCACATCC | RT-qPCR |
| HsSOX2_qPCR_Fw3 | GAGCTTTGCAGGAAGTTTGC | RT-qPCR |
| HsSOX2_qPCR_Rv3 | GCAAGAAGCCTCTCCTTGAA | RT-qPCR |
| HsNANOG_qPCR_Fw4 | AACTCTCCAACATCCTGAACC | RT-qPCR |
| HsNANOG_qPCR_Rv4 | CCTTCTGCGTCACACCATT | RT-qPCR |
| HsTBXT_qPCR_Fw4 | AGGTACCCAACCCTGAGGA | RT-qPCR |
| HsTBXT_qPCR_Rv4 | GCAGGTGAGTTGTCAGAATAGG | RT-qPCR |
| HsTBX6_qPCR_Fw1 | GAACGGCAGAAACTGTAAGAGG | RT-qPCR |
| HsTBX6_qPCR_Rv1 | GTGTGTCTCCGCTCCCATAG | RT-qPCR |
| HsCDX2_qPCR_Fw1 | ATCACCATCCGGAGGAAAG | RT-qPCR |
| HsCDX2_qPCR_Rv1 | TGCGGTTCTGAAACCAGATT | RT-qPCR |
| HsPAX6_qPCR_Fw2 | CGAGATTTCAGAGCCCCATA | RT-qPCR |
| HsPAX6_qPCR_Rv2 | AAGACACCACCGAGCTGATT | RT-qPCR |
| HsOTX2_qPCR_Fw1 | CTCGCCACATCTACTTTGATAGC | RT-qPCR |
| HsOTX2_qPCR_Rv1 | GGTGGACAGGTTCAGAGTCC | RT-qPCR |
| HsNES_qPCR_Fw2 | TCAAGATGTCCCTCAGCCTGGA | RT-qPCR |
| HsNES_qPCR_Rv2 | AAGCTGAGGGAAGTCTTGGAGC | RT-qPCR |
| HsEOMES_qPCR_Fw2 | AAATGGGTGACCTGTGGCAAAGC | RT-qPCR |
| HsEOMES_qPCR_Rv2 | CTCCTGTCTCATCCAGTGGGAA | RT-qPCR |
| HsCER1_qPCR_Fw1 | CTTCTCAGGGGGTCATCTTG | RT-qPCR |
| HsCER1_qPCR_Rv1 | TCCCAAAGCAAAGGTTGTTC | RT-qPCR |
| HsLEFTY1_qPCR_Fw2 | ACCTTGGGGACTATGGAGCT | RT-qPCR |
| HsLEFTY1_qPCR_Rv2 | GCCCACACACTCATAAGCCA | RT-qPCR |
| HsRPL4_qPCR_Fw1 | ACGATACGCCATCTGTTCTGCC | RT-qPCR |
| HsRPL4_qPCR_Rv1 | GGAGCAAAACAGCTTCCTTGGTC | RT-qPCR |
| ONT_RTA_bc01_top | /5PHOS/GGCTTCTTCTTGCTCTTAGGTAGTAGGTTC | DRS |
| ONT_RTA_bc01_bottom | GAGGCGAGCGGTCAATTTTCCTAAGAGCAAGAAGA AGCCTTTTTTTTTT | DRS |
| ONT_RTA_bc02_top | /5PHOS/GTGATTCTCGTCTTTCTGCGTAGTAGGTTC | DRS |
| ONT_RTA_bc02_bottom | GAGGCGAGCGGTCAATTTTCGCAGAAAGACGAGA ATCACTTTTTTTTTT | DRS |
| ONT_RTA_bc03_top | /5PHOS/GTACTTTTCTCTTTGCGCGGTAGTAGGTTC | DRS |
| ONT_RTA_bc03_bottom | GAGGCGAGCGGTCAATTTTCCGCGCAAAGAGAAA AGTACTTTTTTTTTT | DRS |
| ONT_RTA_bc04_top | /5PHOS/GGTCTTCGCTCGGTCTTATTTAGTAGGTTC | DRS |
| ONT_RTA_bc04_bottom | GAGGCGAGCGGTCAATTTTAATAAGACCGAGCGAA GACCTTTTTTTTTT | DRS |

**Table S2:** Sequencing summary of direct RNA-seq from MShef11 hESCs and trilineage differentiated cells. Each replicate sample of hESC and three germ layers were multiplexed into one run, and sequenced to generate 2.0Gb of output. The mean statistics per sample from each run are shown, with the range over the four samples shown in parentheses. N50 indicates the size adjusted median read length, representing the read length for which the 50th percentile base resides.

| | **Run 1** | **Run 2** | **Run 3** |
|---|---|---|---|
| **Total bases /Mb** | 361.5 (297.2–425.4) | 319.4 (203.3–470.0) | 332.6 (207.3–462.2) |
| **18S reads /k reads** | 172.2 (130.5–209.1) | 131.9 (89.6–190.8) | 138.3 (82.5–197.1) |
| **28S reads /k reads** | 45.1 (27.4–58.5) | 47.2 (29.6–74.9) | 38.9 (21.8–63.0) |
| **18S N50 /nt** | 1753 (1676–1818) | 1815 (1812–1817) | 1816 (1813–1818) |
| **28S N50 /nt** | 3107 (1468–4689) | 3517 (2240–4682) | 3679 (2891–4376) |

**Table S3:** Summary of probability thresholds, related to Figure S2A. Using a target sensitivity of 99.5%, probability thresholds were determined in the IVT samples for each primary nucleotide in each rRNA species.

| rRNA | Primary nucleotide | Final threshold | Nucleotides removed (%) | Precision (%) |
|---|---|---|---|---|
| 18S | A | 0.953126 | 24.21135 | 99.63526 |
| 18S | T | 0.996094 | 19.30326 | 99.85464 |
| 18S | C | 0.964845 | 31.40027 | 99.53552 |
| 18S | G | 0.939453 | 10.69514 | 99.71066 |
| 28S | A | 0.976563 | 27.33533 | 99.61441 |
| 28S | T | 0.996094 | 23.95559 | 99.86284 |
| 28S | C | 0.988282 | 38.59012 | 99.59046 |
| 28S | G | 0.941406 | 15.53711 | 99.52708 |
| 5.8S | A | 0.949219 | 19.09199 | 99.53005 |
| 5.8S | T | 0.992188 | 19.84949 | 99.58532 |
| 5.8S | C | 0.957032 | 32.05085 | 99.51769 |
| 5.8S | G | 1 | 9.074344 | 100 |
